# Determinants of Viscoelasticity and Flow Activation Energy in Biomolecular Condensates

**DOI:** 10.1101/2022.12.30.522262

**Authors:** Ibraheem Alshareedah, Anurag Singh, Alexander Quinn, Priya R. Banerjee

## Abstract

The form and function of biomolecular condensates, which are phase-separated intracellular granules of proteins and RNAs, are regulated by their material and dynamical properties. Emerging reports suggest that biomolecular condensates are viscoelastic network fluids, and the primary sequence and structure of the constituent biopolymers govern their bulk fluid phase properties. Here, we employ a multi-parametric approach to dissect the molecular determinants of condensate viscoelasticity by studying a series of condensates formed by engineered multivalent arginine-rich polypeptides and single-stranded DNA. By measuring the terminal relaxation time of the condensate network through optical tweezer-based microrheology and the activation energy of viscous flow through temperature-controlled video particle tracking, we show that condensate viscoelasticity is controlled by two distinct factors − sequence-encoded inter-chain interactions of associative polymers and entropic factors emerging from their intrinsic polymer properties such as the chain length. The biomolecular diffusion in the dense phase shows a strong dependence on the flow activation energy, indicating that the intra-condensate transport properties are primarily reaction-dominant. These results provide a glimpse of the multifaceted control of viscoelasticity and transport properties within biomolecular condensates. Flow activation energy measurement of single and multicomponent condensates by thermo-rheology provides a direct route to quantify inter-chain interactions in the dense phase and dissect the roles of chain entropy and valence in dictating the viscoelastic behavior of biomolecular condensates.

## Introduction

Biomolecular condensates are phase-separated intracellular granules harboring multiple proteins, nucleic acids, and other biomolecules and are ubiquitous in almost all living systems[1, 2]. They have been implicated in key biological processes including stress response[3, 4], gene regulation[5], genome organization and maintenance[6], mitochondrial signaling processes[7], and intracellular storage[8]. Further, aberrant condensates are thought to be involved in disease processes including neurodegenerative disorders and certain types of cancer[9-14]. From an engineering standpoint, biomolecular condensates offer programmable and biocompatible self-assembled soft colloidal structures that have significant potential as artificial organelles, in intracellular cargo delivery and controlled release, and in creating stimuli-responsive artificial cell-like entities[15-20]. Therefore, understanding the fundamental physics of biomolecular condensates, such as their material properties and network structure[21] and how they are linked to the specific features of the component biopolymers, is an important active area of the current research in the field.

Recently, we and others have shown that reconstituted biomolecular condensates are complex fluids with viscoelastic properties that are present in both homotypic condensates formed by a single protein component and heterotypic condensates formed by proteins and nucleic acids[15, 22, 23]. Further, studies in live cells have indicated that the nucleolus, an archetypal protein-nucleic acid condensate that is responsible for rRNA biogenesis and processing, displays viscoelastic behavior[24]. The viscoelastic properties of the nucleolus have been proposed to be an important physical determinant of its function in facilitating the outward flow of processed rRNA[24-27]. Collectively, these recent advances point to unique and complex material properties of biological condensates, which are likely to play key roles in dictating their biological functions. In addition, it has also been suggested that changes in the material properties of some ribonucleoprotein condensates, such as liquid-to-solid transitions, may lead to pathological outcomes[12, 28-30]. Therefore, understanding the origins of biomolecular condensate viscoelasticity will offer key insights into condensate biophysics, biology, and aberrant behaviors in disease processes.

Deciphering the origins of condensate viscoelastic properties is a difficult task due to the complex dependence of these properties on many physicochemical factors including chain length, intermolecular interactions, and the structure of constituent proteins and nucleic acids[26, 31-34]. Previously, we utilized laser tweezer-based microrheology to measure the viscoelastic moduli of a series of designed peptide-RNA condensates and showed their strong dependence on the inter-chain attractions[15]. Additionally, chain entanglement and steric factors such as intermolecular friction in the dense phase can give rise to a dominant viscoelastic behavior in complex fluids[35]. Therefore, it is conceivable that the material properties of biomolecular condensates are governed by both enthalpic contributions (e.g. intermolecular interactions) and entropic effects (e.g. chain entanglement and steric factors). However, the enthalpic and entropic factors are typically convolved and most of the current experimental approaches cannot distinguish whether the observed dynamical properties of a condensate are dominated by intermolecular interactions and/ or by entropic factors.

Motivated by this knowledge gap, here we employ a multiparametric approach to dissect the effect of intermolecular interactions (enthalpy) and chain length (entropy) on the viscoelastic and transport properties of protein-nucleic acid condensates. We utilize our previously designed Arg/Gly-rich (R/G-rich) repeat polypeptides and single-stranded DNA (ssDNA) as model systems that form viscoelastic condensates with a broad range of material properties[15]. Due to the modular design of our peptides and ssDNA components, these synthetic condensates provide a suitable model system to explore the distinct effects of sequence-encoded biomolecular interactions and the effect of polymer chain length on the condensate dynamical properties in a systematic fashion. Firstly, we employ optical tweezer-based microrheology[15] to measure the **viscoelastic shear moduli and the timescale for the network flow** of heterotypic peptide-ssDNA condensates. Next, we perform temperature-controlled video particle tracking (VPT) to probe the **activation energy for the network flow** of these condensates. We show that peptide-ssDNA condensates obey the time-temperature superposition principle[36] and follow the Arrhenius Law of viscosity[37-40], leading to a measurable flow activation energy. We then systematically probe the dependence of viscoelastic moduli, terminal relaxation time, and the flow activation energy on intermolecular interactions, encoded by the polypeptide sequence, and on entropic effects through the ssDNA length variation. We find that while varying intermolecular interaction strength alters the flow activation energy and the condensate viscoelastic properties in a correlative manner, changing the ssDNA length enhances the viscoelasticity of these condensates without any significant change in the flow activation energy. The variation of condensate viscoelasticity with intermolecular interactions and ssDNA length is consistent with a network fluid model[21] which suggests that the viscoelastic properties of condensates are a function of both enthalpic factors (inter-chain interactions) and entropic factors that stem from the polymeric features of the chain[41-44]. On the other hand, a near-constant activation energy as a function of ssDNA length suggests that the flow activation energy is primarily dependent on intermolecular interactions and not on the polymer length in our system. These findings can be rationalized in light of the theory of rate processes as well as recent models of polymer dynamics such as the sticky rouse model[37, 45] and they suggest that flow activation energy is a direct reporter of the strength of intermolecular interactions in the dense phase. We further show that while the polypeptide diffusion in the dense phase is inversely correlated with the flow activation energy (and hence the strength of intermolecular interactions), it is not significantly altered with ssDNA length variation, even when such a variation leads to an order-of-magnitude change in the bulk viscosity of the condensates. These results suggest that bulk mechanical properties are not the only factor governing biomolecular diffusion in the dense phase, stressing the importance of concepts such as reaction-limited diffusion to describe intra-condensate transport properties[46, 47]. Overall, our multifaceted analysis of both material properties and the flow activation energy enables us to dissect the origins of viscoelasticity in biomolecular condensates and its effects on biomolecular transport.

## Results

### Peptide-ssDNA condensates follow the Arrhenius law of viscosity and have a well-defined activation energy of viscous flow

The concept of biomolecular condensates being a network fluid[21] implies at least two measurable quantities: the network reconfiguration timescale, *i*.*e*., the timescale of the network flow, and the activation energy of the network flow. We and others have recently employed laser tweezer-based microrheology to quantify the timescale of network flow[15, 22, 23], however, according to our knowledge, the flow activation energy has not been reported for any biomolecular condensates yet. To probe both of these quantities simultaneously, we first utilized a designed heterotypic condensate system formed by multivalent R/G-rich repeat polypeptide, [RGRGG]_5_, and a 40-nucleotide long ssDNA, dT40. R/G-rich intrinsically disordered peptides (IDPs) have previously been shown to provide a modular platform to dissect the roles of sticker and spacer residues on the phase behavior and material properties of heterotypic IDP-RNA condensates[15, 31]. The sticker-spacer classification of associative polymers is based on the recent works to distinguish amino acids that directly contribute to interchain interactions (stickers) and amino acids that modulate the solvation and structural features of the chain (spacers)[41-44, 48, 49]. In the present context, arginine residues are defined as stickers because they enable nucleic acid binding through a hierarchy of electrostatic, cation-π, and π-π interactions[31, 32, 50]. Employing our previously developed passive microrheology with optical tweezers (pMOT) assay[15, 51], we measured the rheological moduli of peptide-ssDNA condensates formed by mixing [RGRGG]_5_ and dT40 in a buffer containing 25 mM MOPS (pH 7.5), 25 mM NaCl and 20 mM DTT (Fig. 1A, SI Appendix, Fig. S1&S2). We found that [RGRGG]_5_-dT40 condensates exhibit viscoelastic behavior similar to a Maxwell fluid with a terminal relaxation time of ∼20 ms and a terminal viscosity of 3.3 ± 0.2 Pa.s (at T = 27 °C; Fig. 1B). The terminal relaxation time represents the longest relaxation time of the network, while the terminal viscosity is the zero-shear viscosity representing the dominant viscous behavior at long timescales[15]. We next probed the flow activation energy by evaluating the temperature dependence of the terminal viscosity of these condensates. To this end, we utilized temperature-controlled video particle tracking (VPT) to measure the terminal viscosity of [RGRGG]_5_-dT40 condensates as a function of temperature ranging from ∼10-70 °C (Fig. 1C&D, SI Appendix, Fig. S3). This temperature range is well below the upper cloud point temperature of these condensates (>90 °C)[15] (SI Appendix, Fig. S4). We observed that increasing temperature leads to a decrease in the viscosity to ∼0.7 Pa.s at 55 °C. Similarly, decreasing temperature led to an increase in viscosity (∼10 Pa.s at 11 °C) of the same condensates. The viscosity variation with temperature for these condensates can be fitted with an exponential decay function (Fig. 1E and SI Appendix, Fig. S3). This exponential scaling of viscosity with temperature is in accordance with the Arrhenius Law for viscosity[38, 40]

**Figure 1.**
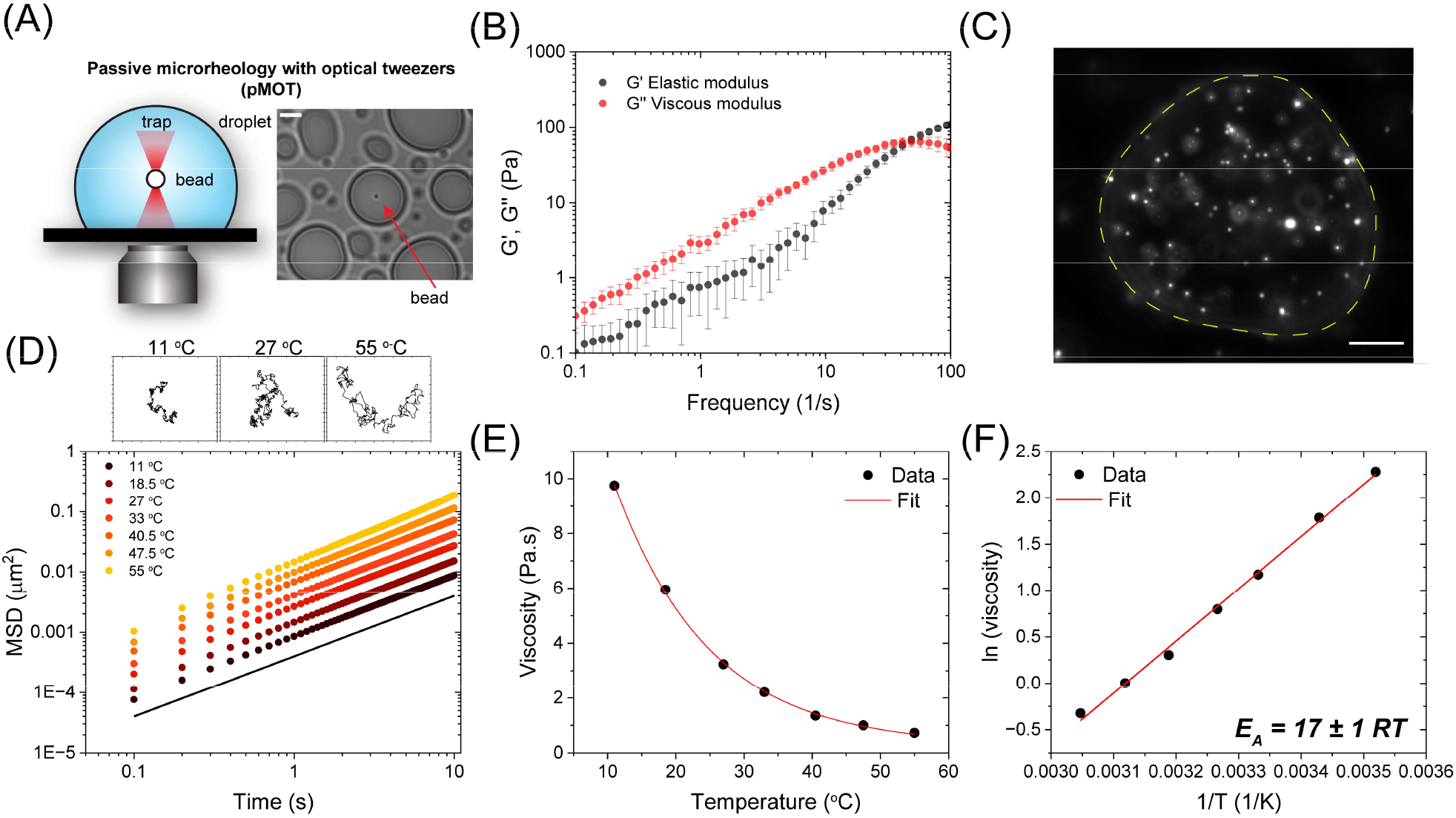
Peptide-ssDNA condensates follow an Arrhenius law of viscosity. **(A)** A scheme showing the experimental setup for the passive microrheology with optical tweezer (pMOT) experiments. **(B)** Average frequency-dependent viscoelastic moduli of [RGRGG]_5_-dT40 condensates (Also see SI Appendix, Fig. S2). **(C)** A representative fluorescence image of 200 nm yellow-green fluorescent beads embedded within an [RGRGG]_5_-dT40 condensate. Scale bar is 10 μm. **(D)** Ensemble-averaged mean squared displacements (MSDs) of the 200 nm beads within [RGRGG]_5_-dT40 condensates at different temperatures. The black line has a slope that corresponds to a diffusivity exponent *α* = 1. The insets show representative particle trajectories at three different temperatures, as indiacted. **(E)** Viscosity of [RGRGG]_5_-dT40 condensates plotted against temperature. The red line is an exponential decay fit using Equation 1 added to a constant. **(F)** Arrhenius plot of viscosity and temperature. The red line represents a linear fit. Activation energy *E*_A_ is calculated from the slope of the line according to Equation 2.

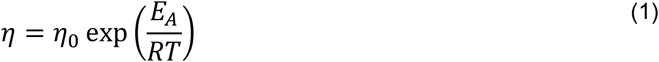

Where *η*_0_ is called the entropic factor or the gas viscosity, *T* is temperature, *R* is the universal gas constant, and *E*_A_ is the activation energy of viscous flow. Accordingly, plotting the natural log of viscosity against the inverse of temperature yields a linear plot with a slope that is equal to *E*_A_/*R*

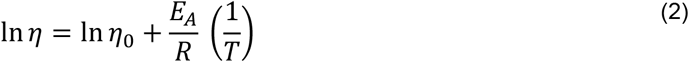

Figure 1F reveals that [RGRGG]_5_-dT40 condensates obey the Arrhenius law (also see SI Appendix, Fig. S3). The resulting activation energy as extracted from the fit was 17 ± 1 RT (at T = 25 °C) which is equivalent to ∼42 kJ/mol. This is comparable to the flow activation energy of 90% Glycerol/Water mixture[52, 53] (SI Appendix, Fig. S5). This indicates that the energy barrier for the condensate network to flow is ∼ 17 times the thermal energy of the system in the standard conditions, as per the theory of rate processes[37]. The observation that the temperature dependence of viscosity for [RGRGG]_5_-dT40 condensate obeys an Arrhenius relation also indicates that these condensates are thermo-rheologically simple fluids within the limits of our experimental temperature range, meaning that they obey the time-temperature superposition principle[54]. This indicates that all the mechanical relaxation modes of the dense phase have a uniform dependence on temperature and are likely to be dominated by the enthalpic forces between the peptide-ssDNA chains.

### The flow activation energy and viscoelastic properties of peptide-ssDNA condensates are governed by sequence-dependent intermolecular interactions

The theory of rate processes by Eyring and others suggests that the flow activation energy of a fluid depends on the enthalpy of interactions amongst the molecular components[37]. For a fluid to flow, the molecules have to rearrange in order to accommodate the exchange of molecules in space (Fig. 2A). Analogous to a chemical reaction, the reconfiguration of a fluid network has an energy barrier that is termed the activation energy of viscous flow[37] (Fig. 2A). When the temperature is increased, the rate of molecules crossing the activation energy barrier increases, leading to a faster flow. Furthermore, when a shear force is applied, the symmetry of the energy barrier is broken such that the rate of molecules crossing the activation barrier in the direction parallel to the flow is higher than those crossing the activation barrier in the opposite direction[37] (Fig. 2A).

**Figure 2.**
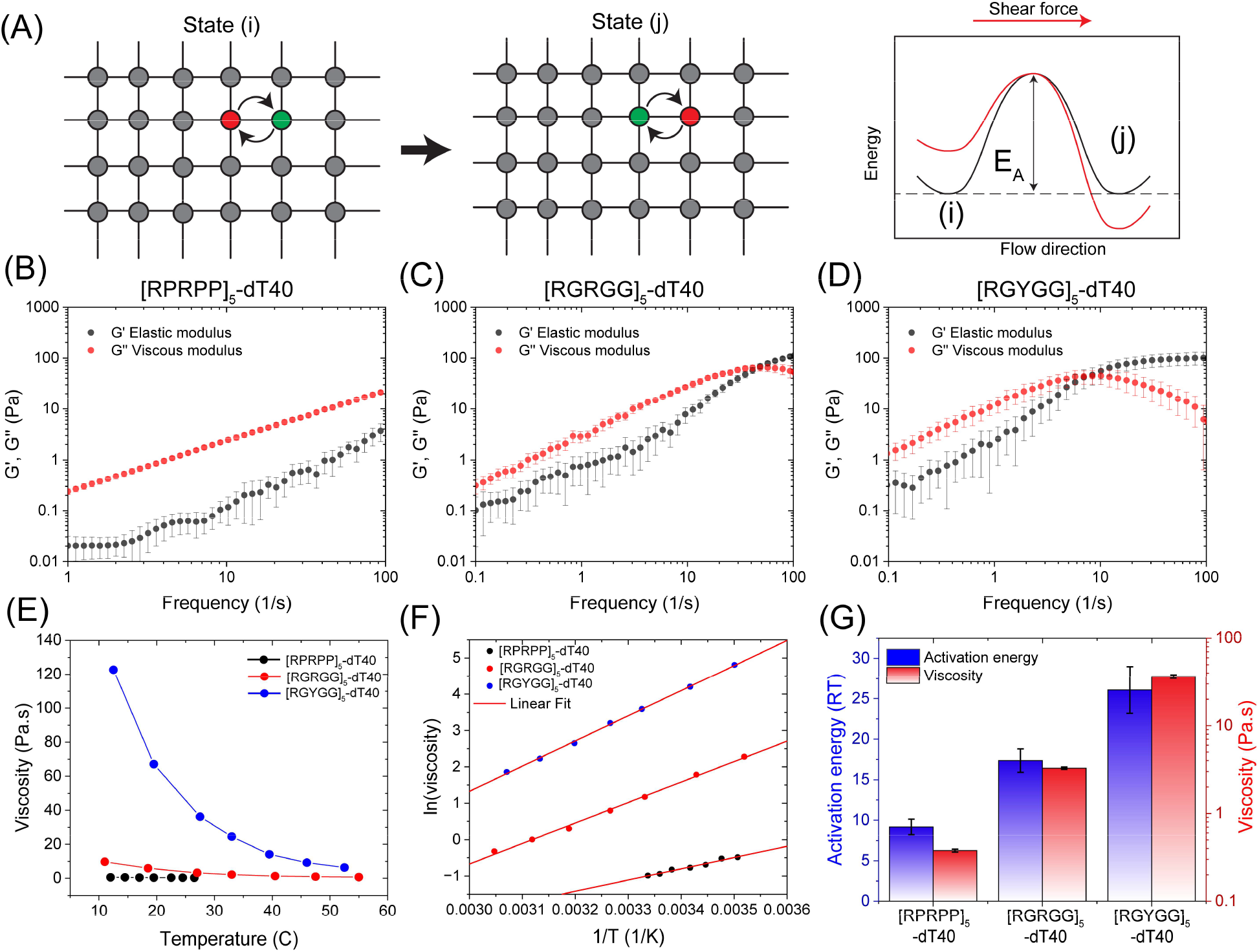
Sequence-encoded intermolecular interactions govern the viscoelastic behavior and flow activation energy of peptide-ssDNA condensates. **(A)** A schematic showing the activation energy as the energy barrier for molecular exchange within the liquid. This barrier is associated with the energy needed to reconfigure the molecules from state (i) to state (j). When a shear force is applied from left to right (red arrow), the barrier becomes asymmetric with the state (j) having lower energy, and hence directional diffusion/flow is favored. **(B-D)** Frequency-dependent average viscoelastic moduli of [RPRPP]_5_, [RGRGG]_5_, and [RGYGG]_5_ condensates with dT40, respectively. Error bars represent the standard deviation estimated from ∼40 data sets. **(E)** Viscosity variation with temperature for peptide-dT40 condensates. The peptides tested are [RGRGG]_5_, [RPRPP]_5_, and [RGYGG]_5_. **(F)** Arrhenius plots of viscosity versus temperature for the three peptide sequences in (B-D). **(G)** Variation of viscosity (at T = 27 °C; red) and flow activation energy (blue) for [RPRPP]_5_, [RGRGG]_5_, and [RGYGG]_5_ condensates with dT40.

Several recent studies have reported that the physical properties of biomolecular condensates are encoded by the constituent protein and nucleic acid primary sequence composition and patterning[18, 48, 55-60]. We therefore asked how the polypeptide sequence features impact the energy barrier for flow? To quantitatively establish a link between the peptide-ssDNA intermolecular interactions and the flow activation energy of the resulting condensates, we employed our three previously characterized nucleic acid-binding repeat peptides with variable sequence composition. Specifically, we previously showed that a variant of [RG<u>R</u>GG]_5_, [RG<u>Y</u>GG]_5_, is a stronger binder to nucleic acids, exhibits higher upper critical solution temperature (UCST) for phase separation, and its condensates with RNA exhibit greater viscoelasticity[15] (SI Appendix, Fig. S4). Contrastingly, a second variant with proline spacers [R<u>P</u>R<u>PP</u>]_5_ displayed the opposite trend, namely, weakened viscoelasticity and lower driving force for phase separation due to weaker RNA binding [15] (SI Appendix, Fig. S4). We posit that if peptide-nucleic acid interactions primarily determine the activation energy of condensate network flow, then condensates formed by these peptide variants will exhibit flow activation energy values that are positively correlated with the relative strength of peptide-nucleic acid binding affinities. We first performed microrheology experiments on individual condensates formed by these three peptides ([RGRGG]_5_, [RGYGG]_5_, and [RPRPP]_5_) with a 40 nucleotide ssDNA, dT40, (Fig. 2B-D) and confirmed that the rank order of condensate viscoelastic properties is [RGYGG]_5_-dT40 > [RGRGG]_5_-dT40 > [RPRPP]-dT40 (Fig. 2B-D), akin to their condensates formed with rU40 RNA[15]. To determine the terminal relaxation time of these condensates, we computed the loss tangent (*G*″/*G*′) (SI Appendix, Fig. S6A). When the loss tangent is less than 1, the elastic modulus dominates the rheological response. We observe that the loss tangent for [RGYGG]_5_-dT40 system crosses 1 at a frequency of ∼10 Hz, indicating a terminal relaxation time of ∼100 ms. For [RGRGG]_5_-dT40, the terminal relaxation time is ∼ 20 ms. Lastly, the [RPRPP]_5_-dT40 condensates have a loss tangent that is greater than 1 over the entire range of experimental frequency, indicating a dominant viscous response. Through VPT experiments, we next measured the terminal viscosity of these condensates at T = 27 °C. Consistent with our laser tweezer-based microrheology experiments, we find that [RPRPP]_5_-dT40 condensates have the lowest viscosity of 0.38 ± 0.02 Pa.s, which is followed by [RGRGG]-dT40 condensates (3.28 ± 0.09 Pa.s), whereas the [RGYGG]_5_-dT40 condensates registered the highest viscosity (37 ± 2 Pa.s) (SI Appendix, Fig. S6B). These results are consistent with our previous report and they reaffirm the dependence of the condensate rheological response at the mesoscale on the peptide-ssDNA interactions at the molecular scale[15].

Next, to measure the flow activation energy of these peptide-ssDNA condensates, we performed temperature-controlled VPT experiments (Fig. 2E-G and SI Appendix, Fig. S6B-F). We find that [RGYGG]_5_-dT40 condensates have an activation energy of 26 ± 3 RT, which is significantly greater than [RGRGG]_5_-dT40 condensates that have an activation energy of 17 ± 1 RT (Fig. 2E-G and SI Appendix, Fig. S6). Further, [RPRPP]_5_-dT40 condensates featuring weakened nucleic acid binding exhibited substantially lower activation energy (9 ± 1 RT; Fig. 2E-G and SI Appendix, Fig. S6). These results reveal a direct correlation between the strength of peptide-ssDNA interactions and the activation energy for the network flow of their respective condensates. Therefore, the strength of intermolecular interactions governs, at least in part, the flow activation energy as well as the viscoelasticity and terminal viscosity of peptide-ssDNA condensates (Fig. 2 and SI Appendix, Fig. S6).

### DNA length alters the viscoelasticity of peptide-ssDNA condensates

In addition to the strength of inter-chain interactions, the viscoelastic response of complex fluids can also be tuned by the length of associative polymer chains[61]. We tested this idea by probing the effect of DNA length on the rheology of peptide-ssDNA condensates. We utilized poly(dT) sequences featuring different numbers of nucleotides ranging from 20 to 200 nucleotides (dT20, dT40, dT90, and dT200). Microrheology experiments on condensates formed by [RGRGG]_5_ with ssDNA of variable length at identical mass concentrations (5 mg/ml peptide and 5 mg/ml ssDNA) showed an increased viscoelastic response with increasing DNA length. For instance, [RGRGG]_5_- dT20 condensates showed a dominant viscous behavior across the experimentally accessible frequency range, with a terminal relaxation time of ∼ 14 ms (Fig. 3A&F) and a terminal viscosity of 1.3 ± 0.1 Pa.s (at 27 °C; Fig. 3G, and SI Appendix, Fig. S7). Increasing the DNA length to 40 nucleotides led to a higher viscosity (3.28 ± 0.09 Pa.s) and a longer terminal relaxation time (∼23 ms; Fig. 3B&E-G). Further lengthening of the DNA chain to 90 nucleotides (dT90) led to an increase in viscosity to 11 ± 2 Pa.s and an increase in relaxation time to ∼100 ms (Fig. 3C&E-G). Finally, increasing the DNA length to 200 nucleotides (dT200) led to a stronger viscoelastic behavior with a viscosity of 23 ± 2 Pa.s and a relaxation time of ∼200 ms, which is more than an order of magnitude higher as compared to the condensates formed by dT20 (Fig. 3D&E-G). These results establish an important role of ssDNA length in dictating the viscoelastic properties of these condensates (Fig 3 and SI Appendix, Fig. S7).

**Figure 3.**
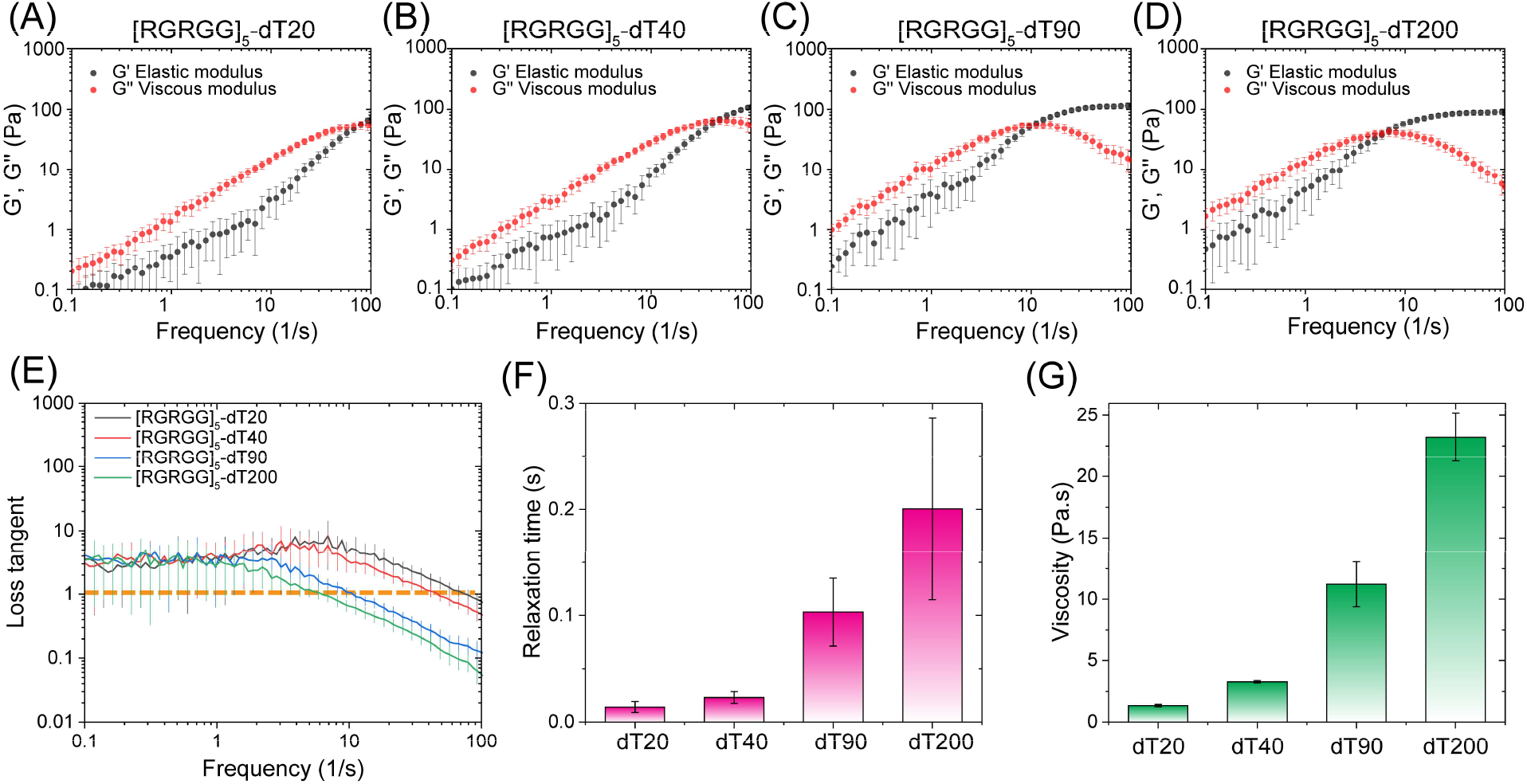
ssDNA length alters the viscoelasticity of peptide-ssDNA condensates. **(A-D)** Average frequency-dependent viscoelastic moduli of [RGRGG]_5_-dT*n* condensates, where *n* is 20, 40, 90, and 200, respectively. **(E)** Loss tangent (*G*″>/*G*′) of four peptide-DNA condensates in (A-D) showing distinct terminal relaxation times with DNA length variation. **(F)** Terminal relaxation times of [RGRGG]_5_-dT*n* condensates corresponding to (A-D). **(G)** Viscosity variation with ssDNA length for [RGRGG]_5_-dT*n* condensates (*n* = 20, 40, 90, and 200) as measured by video particle tracking (VPT) at T= 27 °C.

### DNA chain length does not impact the flow activation energy of peptide-DNA condensates

So far, our results suggest that the dynamical properties of peptide-ssDNA condensates are tunable through two independent factors: (a) varying the ssDNA-peptide interactions (enthalpic driving force), and (b) ssDNA length. To further probe the impact of the polymer length on the dynamical properties of these condensates, we measured their flow activation energy as a function of ssDNA length. Although the condensate viscosity increased by an order of magnitude with increasing ssDNA length from 20 nucleotides (20-nt) to 200 nucleotides (200-nt), temperature-controlled VPT measurements reveal similar exponential decays of condensate viscosity with temperature (Fig. 4A and SI Appendix, Fig S8). Intriguingly, the Arrhenius plots for peptide-ssDNA condensates showed similar slopes irrespective of the ssDNA length (20, 40, 90, and 200-nt), indicating a constant activation energy (Fig. 4B and SI Appendix, Fig S8). This means that the flow activation energy (Equation 2) of these condensates does not change with increasing DNA length (from 20-nt to 200-nt) despite an order of magnitude change in condensate viscoelasticity (Fig. 4C).

**Figure 4.**
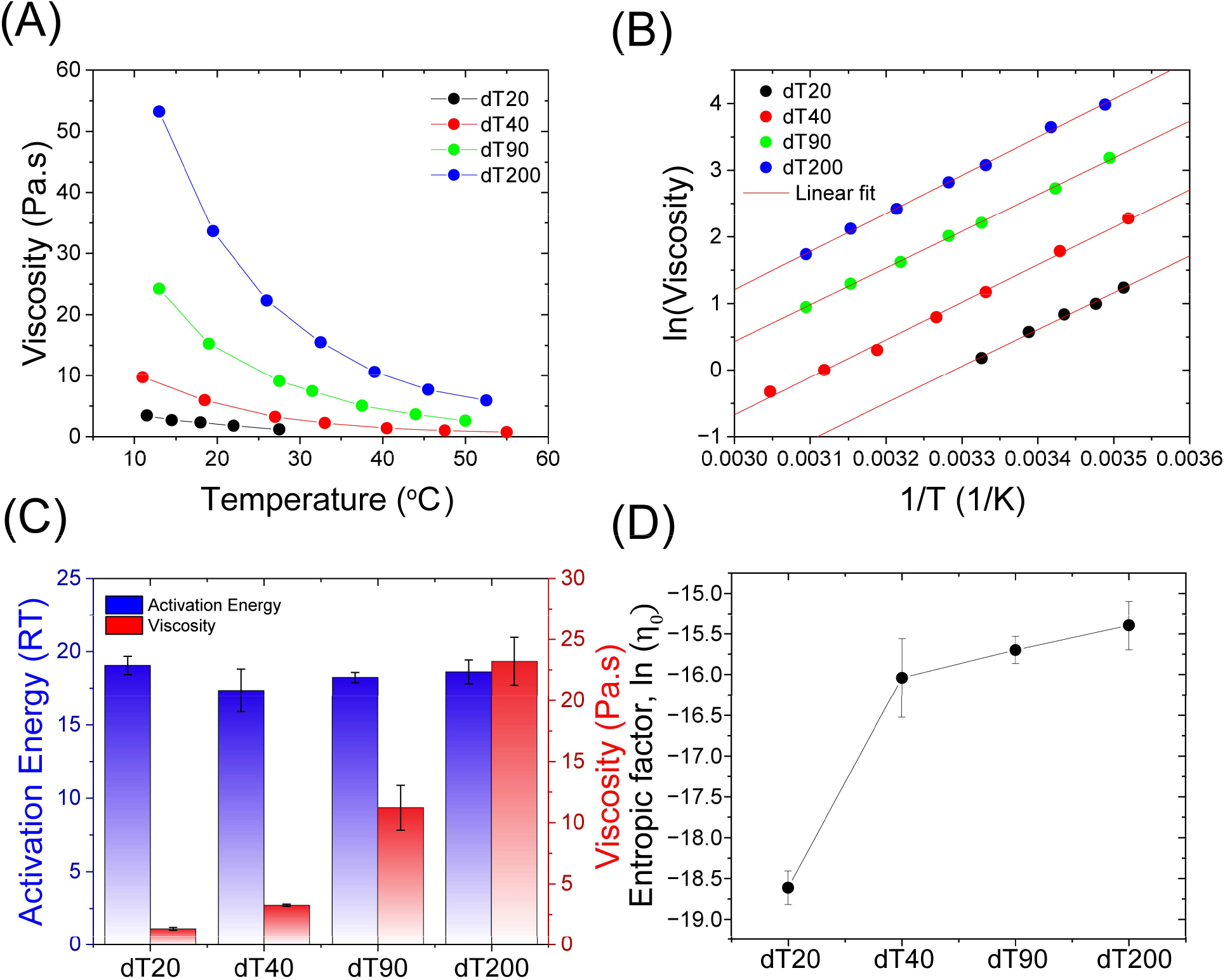
ssDNA length does not affect the flow activation energy of peptide-ssDNA condensates. **(A)** Viscosity variation with temperature for [RGRGG]_5_-dT*n* condensates where *n* ranges from 20 to 200. **(B)** Arrhenius plots for [RGRGG]_5_-dT*n* condensates. Note the similar slopes for all DNA lengths. Red lines are linear fits to the data. **(C)** Flow activation energy and viscosity (at 27 °C) of [RGRGG]_5_-dT*n* condensates. **(D)** The extrapolated entropic factor of peptide-DNA condensates from the data shown in (B). Activation energy *E*_A_ and the entropic factor is calculated from the slope and intercept of the linear fit according to Equation 2. Also see SI Appendix, Figure S9.

According to Eyring’s transition state theory of diffusion, the flow activation energy of the system depends on the enthalpic interactions between the polymers chains[37]

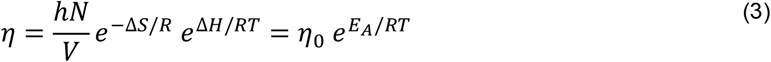

Where *h* is Planck’s constant, *N* is the number of molecules, *V* is the volume, and *R* is the universal gas constant. Δ*H* and Δ*S* are the entropy and enthalpy changes when two neighboring molecules switch positions within the fluid. This relation indicates that the activation energy, *i*.*e*., the energy barrier of reconfiguring the fluid network to achieve the exchange of molecules between the two neighboring positions, is predominantly determined by the intermolecular interactions between the polymer chains (Fig. 2A). Therefore, according to Equation 3, our results suggest that changing ssDNA length does not alter the extent of enthalpic forces within the condensate network. What then could be the origin of the enhanced viscoelasticity of these condensates upon increasing ssDNA length (Fig. 3)? A closer inspection of the Arrhenius plots of these peptide-ssDNA condensates reveals that the pre-exponential factor increases with the length of the ssDNA (Fig. 4D). This pre-exponential factor (Equations 1&3) is often termed the entropic factor since it is thought to contain information on the entropy change that is associated with the flow (Equation 3)[39, 62]. These data suggest that the changes in the material properties of condensates upon ssDNA length variation are likely due to entropic factors that stem from the increased ssDNA chain length. Taken together with the results reported in Figures 2&3, we conclude that the quantification of the flow activation energy can provide direct insights into the distinct roles of intermolecular interactions and polymer chain length underlying the dynamical properties of peptide-ssDNA condensates.

### The sticky Rouse model explains multiple pathways to regulate condensate viscoelasticity

To conceptualize the variation of flow activation energy and viscoelasticity with intermolecular interactions and ssDNA chain length, we consider the sticky Rouse model for associative polymers such as oppositely charged polyelectrolytes[45]. Our experimental results reported above can be summarized in two statements: First, we observe that stronger intermolecular interactions enhance the viscoelastic response of peptide-DNA condensates as well as their flow activation energy. Second, increasing ssDNA length enhances the viscoelastic behavior of these condensates without altering the flow activation energy. According to the sticky polymer dynamics model[45], a polymer chain undergoes phase separation through associative intermolecular interactions among the binding sites on the chain (Fig. 5A). These binding sites are termed stickers following the seminal work by Rubinstein and Semenov on thermo-reversible gelation of associative polymers and the subsequent works of Pappu and colleagues [41-44, 63, 64]. The stickers in peptide-DNA mixtures can be considered as the R*x*R/Y*xx* motifs (*x* = Gly or Pro) on the peptide and the phosphate backbone and nucleobases on the ssDNA chain[15], which can interact via electrostatic and cation−π interactions[31, 32]. The association time between two stickers can be written as[45, 65]

**Figure 5.**
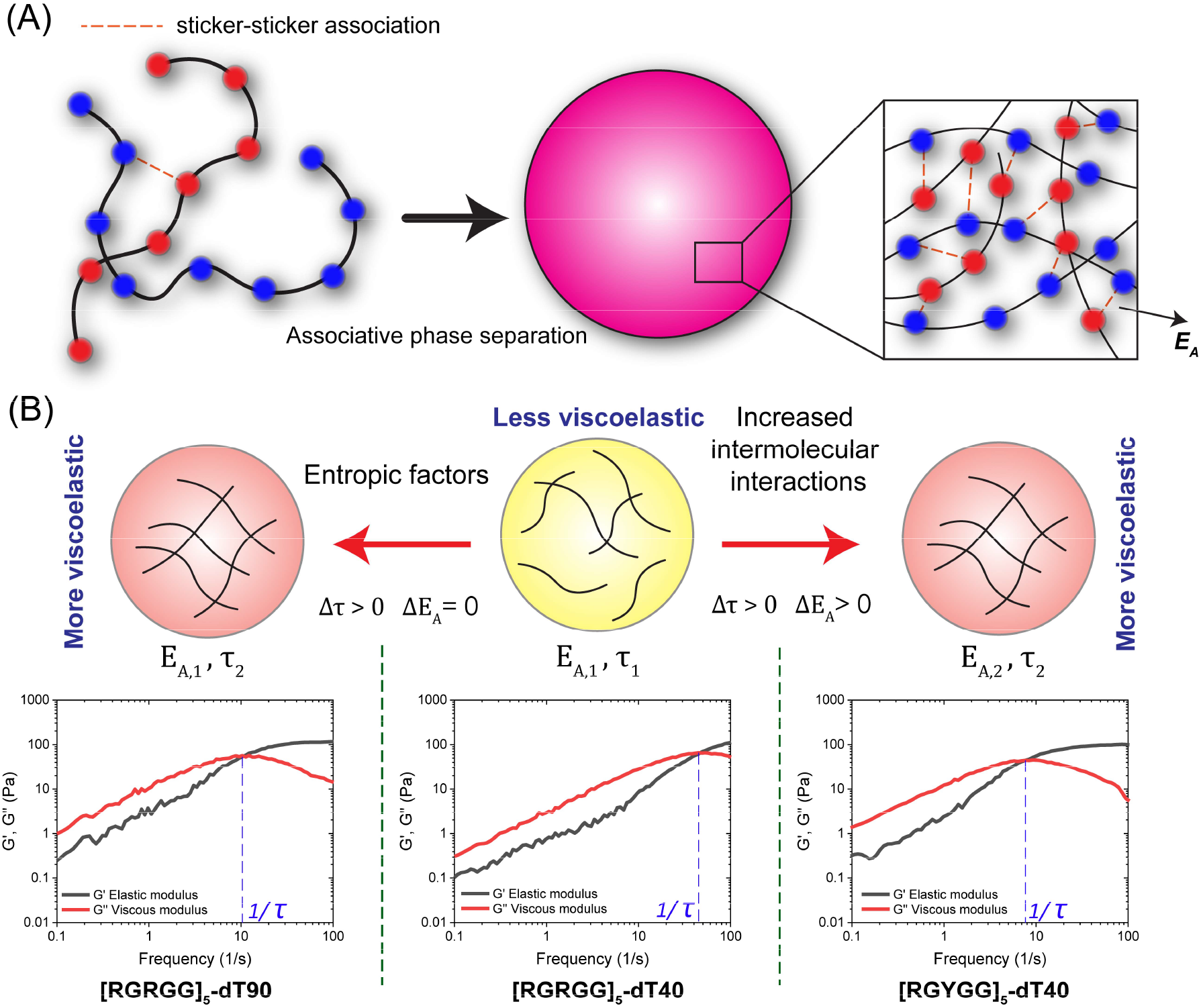
Alternative routes to regulate condensate viscoelasticity. **(A)** A schematic of peptide-ssDNA mixtures represented by chains with complementary stickers that undergo heterotypic phase separation via sticker-sticker association. **(B)** Two thermodynamic pathways for condensate viscoelasticity that are distinguishable by thermo-mechanical analysis. Enhanced viscoelasticity can arise from increasing intermolecular interactions which changes the activation energy of the condensates. Alternatively, enhanced viscoelastic behavior can arise from increasing chain length and the number of stickers, which does not alter the activation energy. Plots of average viscoelastic moduli are shown in the bottom panels as examples of two condensate systems that have similar viscoelastic properties but are designed through different pathways: [RGYGG]_5_-dT40 condensates exhibit stronger viscoelastic response due to stronger intermolecular interactions while [RGRGG]_5_-dT90 condensates achieve similar viscoelastic behavior through longer ssDNA chain length.

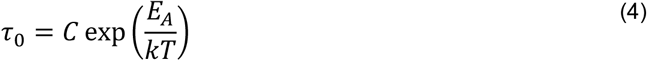

Where *C* is the association time in absence of any attractive interactions and *E*_A_ is the energy of the associative bond. *E*_A_ can be also regarded as the activation energy of the association, which is related to the flow activation energy that we measured in our experiments. The relaxation modes of the material can be written as[45]

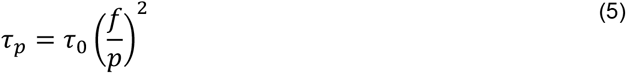

Where *p* is the number of bound stickers per chain and *f* is the total number of stickers (bound + unbound) per chain. The terminal relaxation time of the condensate is equivalent to the case when *p* = 1, wherein the chain becomes completely free to diffuse within the condensate. Hence, the terminal relaxation time *τ*_*R*_ can be written as

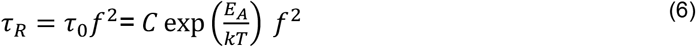

From this relation, it is clear that the enhancement of condensate viscoelasticity can originate from two distinct pathways (Fig. 5B). Longer *τ*_*R*_ can be a result of higher *τ*_0_ and/or larger *f*. Higher *τ*_0_ indicates stronger inter-chain interactions and corresponds to a higher activation energy *E*_A_. Hence, according to Equations 4 and 6, increasing inter-molecular interactions will result in a larger *E*_A_ and *τ*_*R*_. On the other hand, increasing the valence (or length) of DNA translates to an increased number of stickers per chain *f*, which can independently increase the terminal relaxation time *τ*_*R*_ without increasing the association lifetime *τ*_0_ or the activation energy *E*_A_. Alternatively, if one increases the size of the chain without increasing the number of stickers, one would also expect an increase in *τ*_*R*_ [66, 67]. Such an increase in *τ*_*R*_ with increasing chain length can be explained on the basis of the constitutive equation of the Maxwell model[68]. A Maxwell fluid is described using the mechanical analogs of elasticity (spring) and viscosity (dashpot). For a spring, the relation between the stress *σ* and the strain *γ* is

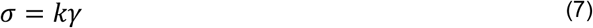

Where *k* is the spring constant. This represents the physical cross-links that occur within the condensate network through intermolecular interactions. For the dashpot, the stress-strain relation is given by

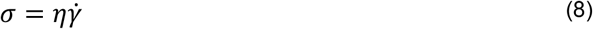

Where *η* is the fluid viscosity. When both elements are connected in series, the constitutive equation can be written as[68]

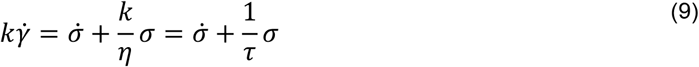

Where *k* is the spring constant, *γ* is the strain, *σ* is the stress, *k* is the spring constant, *η* is the viscosity, and 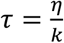 is the relaxation time that is analogous to the terminal relaxation time measured using our laser tweezer-based microrheology experiments of the peptide-ssDNA condensates. Previous experiments and theories have established that with increasing polymer chain length, the viscosity *η* of a polymer solution increases[66, 67, 69, 70]. Consequently, a higher *η* leads to a longer relaxation time *τ*_*R*_ (since *τ*_*R*_ = *η*/*k*).

Therefore, while enhanced viscoelasticity can arise from stronger intermolecular interactions, other factors such as a higher number of stickers and/or polymer chain length can independently impact the condensate viscoelastic behavior. Laser tweezer-based microrheology in conjunction with activation energy measurements can distinguish between these pathways based on their distinct effects on the flow activation energy (Fig. 5B).

### Polypeptide diffusion in the dense phase is strongly correlated with the flow activation energy

A key relevance of quantifying the material properties of biomolecular condensates lies in understanding how macromolecular transport is regulated in the dense phase. It is generally accepted that biomolecular diffusion reports on the material state of the condensates, with slower diffusion indicating higher viscosity of the dense phase. The perceived correspondence between molecular diffusion and material properties is one of the core principles used to infer the material states of condensates using Fluorescence Recovery after Photobleaching (FRAP)[71]. This correlation is based upon the assumption that the Stokes-Einstein equation, or some version of it, is applicable to model biomolecular diffusion within these condensates. However, transport properties of associative biopolymers, in theory, can be dominated by a reaction-diffusion mechanism rather than pure diffusion within a biomolecular condensate, especially when one considers the reversible association and dissociation of the chain with the condensate viscoelastic network[71]. In addition, the viscoelastic network structure can result in a length scale-dependent molecular transport in the dense phase[72, 73]. Our microrheology results revealing distinct thermodynamic forces to tune condensate material properties prompted us to ask how these forces, viz. intermolecular interactions (enthalpic) and polymer chain length (entropic), regulate biomolecular diffusion in the dense phase.

To probe the correlation between the viscoelastic properties and biomolecular transport, we chose three condensate systems. Our reference condensate is [RGRGG]_5_-dT40, which has a bulk viscosity of 3.28 ± 0.09 Pa.s and an activation energy of 17 ± 1 RT. The first variant system is [RGYGG]_5_-dT40 condensate, which has a ∼10-fold higher viscosity (37 ± 2 Pa.s) and a higher activation energy of 26 ± 3 RT. This system is a representative of an altered intermolecular interaction pathway (Fig. 5B). The second variant system is [RGRGG]_5_-dT200, which has a significantly higher viscosity of 23 ± 2 Pa.s than the reference condensate but a similar activation energy of 18.6 ± 0.8 RT. This variant represents an entropic route to enhance condensate viscoelasticity (length variation, Fig. 5B). To estimate the diffusion timescale of polypeptides, we performed fluorescence recovery after photobleaching (FRAP) experiments using fluorescently labeled polypeptides under identical experimental conditions (Fig. 6A&B; *see* SI Appendix, Materials and Methods). In all condensates tested, near-complete FRAP recovery is observed. We calculated the apparent diffusion timescale (*τ*_*D*_) of the peptides as the FRAP recovery half-time normalized with respect to the radius of the bleaching area (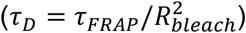) [74-76]. We find that [RGRGG]_5_-dT40 condensates have a diffusion timescale of *τ*_*D*_ = 36 ± 6 s/μm^2^ for the peptide ([RGRGG]_5_). For [RGYGG]_5_-dT40 condensates, the peptide dynamics are significantly slowed down, with [RGYGG]_5_ peptide diffusion timescale of *τ*_*D*_ = 190 ± 30 s/μm^2^ (Fig. 6C&D). This is not surprising given that [RGYGG]_5_-dT40 condensates have a higher bulk viscosity than [RGRGG]_5_-dT40. Contrastingly, when changing the DNA length ([RGRGG]_5_-dT200), the [RGRGG]_5_ peptide diffusion was unaltered, as evidenced by an almost identical FRAP recovery pattern and a similar diffusion timescale (59 ± 6 s/μm^2^, Fig. 6C&D). In fact, the FRAP recovery traces of [RGRGG]_5_ peptide appears almost insensitive to varying ssDNA length from 20-nt to 200-nt (Fig. 6C&D), even though the bulk viscosity changes by almost an order of magnitude (Figs. 3&4) with such variation (*τ*_*D*_ is 56±4, 36±6, 65±3, and 59±6 s/μm^2^ for dT20, dT40, dT90, and dT200 condensates, respectively). These results indicate that bulk viscosity does not govern polypeptide diffusion within these condensates (Fig. 6D top panel), which would be expected if the Stokes-Einstein relation and Fick’s law of diffusion hold true for these condensates. Instead, our results demonstrate that there is a strong correlation between the flow activation energy and the peptide diffusion within peptide-ssDNA condensates (Fig. 6D bottom panel). This observation indicates that the diffusion dynamics of peptides are likely to be governed by a reaction-dominant mechanism where the enthalpy of peptide-DNA interactions regulate macromolecular transport in the dense phase.

**Figure 6.**
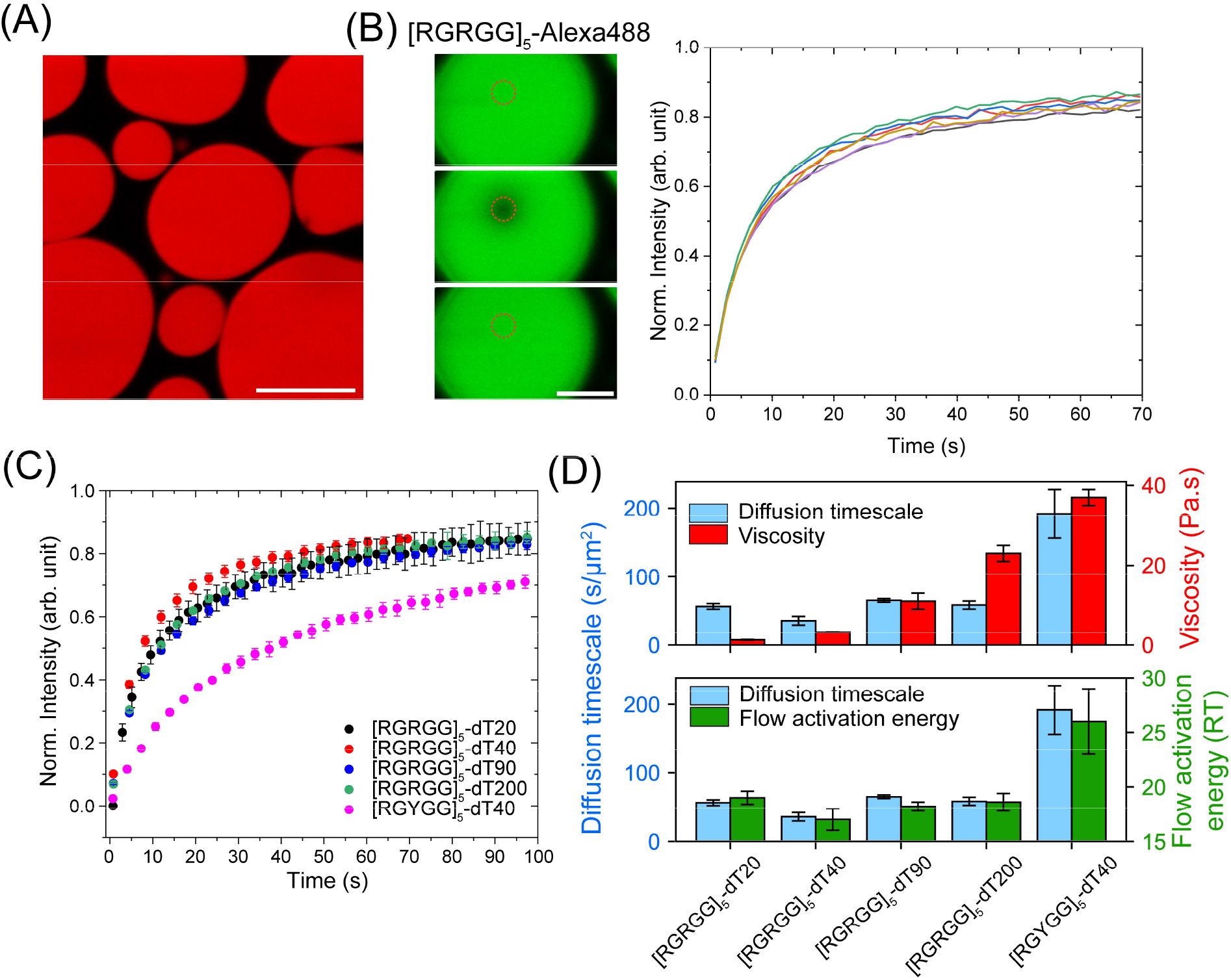
Polypeptide diffusion within peptide-ssDNA condensates scales with flow activation energy but not with the bulk viscosity of the dense phase. **(A)** Representative fluorescent image of [RGRGG]_5_-dT40 condensates visualized by Cy5-labeled DNA. Scale bar is 10 μm. **(B)** Fluorescence recovery after photobleaching for the Alexa488-labeled [RGRGG]_5_ peptide in [RGRGG]_5_-dT40 condensates. Intensity traces from six different condensates are shown. **(C)** Average FRAP recovery intensity trace for Alexa488-labeled peptide in condensates formed by [RGRGG]_5_ and dT20, dT40, dT90, and dT200, as well as condensates formed by [RGYGG]_5_ and dT40, respectively. The value of the intensity is an average of 4-6 trials and the error is the standard deviation of the same. **(D)** Diffusion timescale of Alexa488-labeled peptides in various peptide-DNA condensates as calculated from (C). Also plotted are the viscosity of the condensates (red, top panel) at T=27 °C and the flow activation energy of the same condensates (green, bottom panel). All samples were prepared at 5 mg/ml peptide and 5 mg/ml ssDNA in a buffer containing 25 mM MOPS (pH 7.5), 25 mM NaCl, and 20 mM DTT.

## Discussion

The material properties of biomolecular condensates have garnered significant attention due to their roles in controlling the condensate function in the cell[77]. Viscoelastic behavior, marked by the presence of a dominant elastic response at short timescales but a dominant viscous response at longer timescales, is a hallmark of most types of condensates[15, 22-24, 42, 78]. This is typically due to the polymeric nature of biomolecules forming these condensates and the percolation transition associated with phase separation in these systems[21, 79]. Previously, we have demonstrated a programmable viscoelastic behavior of heterotypic condensates formed by RNA-binding repeat polypeptides and RNA by tuning intermolecular interactions through peptide sequence design[15]. Simultaneously, several pieces of evidence emerged demonstrating viscoelasticity as a common trait in heterotypic and homotypic condensate systems[22, 23, 78]. In a recent report, RNA entanglement was proposed to be the cause of the viscoelastic behavior of the nucleolus [24]. These advancements point to a complex interplay between intermolecular interactions and entropic factors stemming from the polymer effects of disordered protein and/or nucleic acid chains in governing the viscoelastic behavior of biomolecular condensates[24]. Therefore, dissecting the distinct roles of enthalpy and entropy in determining condensate viscoelasticity is essential for understanding the condensate internal network structure as well as for devising suitable strategies to tune its material properties. In this work, we have introduced a multiparametric approach to probe the origin of condensate viscoelasticity by combining microrheology measurements with thermo-rheological analysis in a series of designed peptide-ssDNA condensates.

Firstly, we showed that peptide-DNA condensates obey the Arrhenius law of viscosity, which indicates that all mechanical relaxation modes within the condensate have a uniform temperature dependence[38, 40, 80]. In other words, all physical cross-links within the condensate network respond in a similar fashion when changing temperature, even if they form and disassociate on different timescales. This led us to quantify the condensate flow activation energy, which ranges from 9−26 *RT*. These values are within the same range as previously reported flow activation energies for synthetic complex coacervates using bulk rheology[45, 65, 81-83]. Altering intermolecular interactions through peptide sequence variations lead to a correlative change in the flow activation energy, indicating that the energy barrier of the condensate network flow is primarily governed by the interaction strength of the associative polymers. Further, an increase in intermolecular interactions leads to an enhanced viscoelastic behavior as evident from increased viscosity and terminal relaxation time of the condensates. In contrast, changing the ssDNA length from 20-nt to 200-nt did not alter the flow activation energy of the condensates. However, significant changes in the viscoelastic behavior of the condensate were observed upon such length variation. We propose that the origin of altered condensate viscoelasticity upon ssDNA length variation is entropic in nature. Simply put, due to the presence of larger DNA chains within the condensates, the viscous drag force is increased and translational motion of the chains are slowed down. Due to such a decrease in chain mobility, the elastic regime becomes dominant over longer timescales. Further, larger chains with increased valence entail a higher number of physical crosslinks per chain needed to be relaxed in order for the chain to move[83]. These effects can result in an increase in the pre-exponential entropic factor upon lengthening the DNA (Fig. 4D). Collectively, these results suggest that quantification of the condensate activation energy through thermo-rheology can provide direct insight into the inter-chain interactions in the dense phase and the effects of chain entropy and valence in dictating the viscoelastic behavior of biomolecular condensates.

The simultaneous quantification of flow activation energy and viscoelasticity in our designed condensates allowed us to probe the distinct roles of intermolecular interactions and chain length on biomolecular transport in the dense phase. Unexpectedly, we observed that the translational mobility of molecules within condensates does not scale with the bulk viscosity (Fig. 6D). Rather, diffusion measurements in the dense phase show that the peptide mobility scales with the flow activation energy of the condensate (Fig. 6D). This indicates that the translational motion of the peptides within the condensates is reaction-limited rather than purely diffusive[46, 47]. These results highlight the utility of our combinatorial approach of measuring flow activation energy and viscosity in determining the mechanism of biomolecular transport within condensates. Our approach allows for a mechanistic explanation of the distinct dynamics of biomolecules within heterotypic biomolecular condensates that were reported previously[84, 85]. For example, Keenen et al. studied the condensation of heterochromatin protein 1α (HP1α) with dsDNA[84]. Using FRAP experiments, it was reported that HP1α exhibits identical dynamics within HP1α- dsDNA condensates irrespective of the size of the dsDNA[84]. Based on our results reported here, we expect that increasing the dsDNA length enhances the viscosity of the condensates while leaving the flow activation energy of the condensates unchanged. The insensitivity of HP1α dynamics to the DNA chain length indicates that the protein diffusion within these condensates is primarily governed by a reaction-dominant mechanism. Importantly, these observations, in conjunction with the results reported here, suggest that probing the diffusivity dynamics of macromolecules does not necessarily reflect the material properties of biomolecular condensates, especially if the translational mobility of the macromolecule is dominated by the kinetics of binding and unbinding with the condensate network.

In a broader sense, heterotypic biomolecular condensates contain different types of macromolecules which come together through a hierarchy of intermolecular interactions[86]. The concentrations of macromolecules in the dense phase are thought to depend on this interaction hierarchy, adding a layer of sequence specificity to the compositional control of biomolecular condensates[87, 88]. Our results here lead us to propose another layer of specificity that pertains to the control of molecular dynamics. Within a viscoelastic biomolecular condensate, macromolecules can exhibit distinct diffusivity dynamics depending on their interactions with the condensate viscoelastic network. Recent theoretical work discussed the possibility of a noise-buffering mechanism within cells through phase separation, which regulates the protein concentrations within condensates even when the protein expression levels change throughout the cell cycle[8, 89]. As shown in this work and previous reports, the material properties of heterotypic condensates are sensitively dependent on several factors including sequence and chain length of component biomolecules, ionic strength, and pH among other factors[15, 22-24, 45, 78]. However, we observe that the diffusion of polypeptides (or HP1α in Keenen et al.[84]) within protein-NA condensates can be insensitive to changes in condensate bulk material properties. Based on these results, we speculate that a similar buffering of the mobility rates of macromolecules within condensates can be achieved through a reaction-dominant mechanism. In such cases, even if the condensate material properties change due to external or internal factors, the dynamics of specific components such as our polypeptides can remain unaltered.

In summary, we combine optical tweezer-based microrheology and temperature-controlled fluorescence microscopy to present a thermo-rheological analysis of a series of viscoelastic peptide-ssDNA condensates. Our multiparametric approach allows us to directly probe the contributions of enthalpic interactions and entropic effects on the condensate material properties. Our results shed light on the altered viscoelastic behavior of condensates upon sequence and length variation of constituent biopolymers as well as their impact on the biomolecular transport properties. Importantly, our findings have implications on the way macromolecular diffusion within condensates can be interpreted. We envision that our experimental approach of quantifying condensate viscoelasticity along with flow activation energy will enable precise comparisons of biomolecular condensates that feature component biomolecules with distinct sequences, sizes, and structures. The experimental approach we introduced here provides a deeper understanding of the material properties of biomolecular condensates and their effects on molecular dynamics beyond the simple characterization of the condensate viscoelastic moduli. Such understanding can ultimately enable rational strategies to design and manipulate the material and transport properties of biomolecular condensates.

## Acknowledgements

The authors gratefully acknowledge informative discussions with Rohit V. Pappu and Tanja Mittag and their group members. P.R.B acknowledges funding from the National Institute of General Medical Sciences (NIGMS) of the National Institutes of Health for financial support (R35 GM138186). P.R.B. gratefully acknowledges additional funding support through the St. Jude Research Collaborative on Biophysics of RNP granules.

## Author Contributions

I.A. and P.R.B. conceived the idea and designed this study and the experiments. I. A. developed procedures for experimental data analysis. I.A. and A.S. curated the data with help from A.Q. I.A., A.S., and A.Q. analyzed the data. I.A. and P.R.B. wrote the manuscript. All authors contributed to revising the manuscript.

