## Supplementary Information for "Determinants of Viscoelasticity and Flow Activation Energy in Biomolecular Condensates"

Ibraheem Alshareedah[§]*, Anurag Singh[§], Alexander Quinn, and Priya R. Banerjee*
Department of Physics, University at Buffalo, Buffalo NY 14260

[§] These authors contributed equally

* Correspondence should be addressed to:
P. R. B.:; I. A.:

**This PDF file includes:**

Supplementary text
Supplementary Table S1
Supplementary Figures S1 to S9
SI References

### Materials & Methods

**Materials.** ssDNA oligos of various lengths (20, 40, 90, and 200) were purchased from Integrated DNA Technologies (NJ, USA). The dry stocks were reconstituted in RNase-free water. The reconstituted solutions were centrifuged at 23000xg for 2 minutes to remove any particles and the supernatant was extracted. DNA concentration was measured subsequently using a NanoDrop 1C™ spectrophotometer. The DNA stocks were then aliquoted and stored at -20 °C for further use. The peptides $[RGRGG]_5$, $[RGYGG]_5$, and $[RPRPP]_5$ were synthesized by Genscript inc, USA., and were reconstituted in RNase-free water containing 50 mM DTT (dithiothreitol, ThermoFisher Scientific). The purity of the peptides was higher than 90% as per manufacturer specifications. All the peptide sequences contain a C-terminal cysteine that is used for site-specific labeling with fluorescence dyes. Yellow-green carboxylate-functionalized polystyrene microspheres (diameter 0.2 and 1 µm) were purchased from Invitrogen. Cy5-labeled dT40 (IDT) was used for fluorescence imaging of the condensates.

**Sample preparation.** Peptide-ssDNA condensates were prepared by mixing appropriate amounts from concentrated stock solutions of the peptide and DNA to a final concentration of 5 mg/ml peptide and 5 mg/ml ssDNA. Thus, the mass ratio of the DNA to the peptide is 1:1. This ratio was chosen based on turbidity measurements suggesting that the 1:1 ratio has maximal phase separation for all the systems tested (Fig. S1). The sample buffer contained 25 mM MOPS (pH 7.5), 25 mM NaCl, and 20 mM DTT. MOPS buffer was chosen because it has less pH dependence on temperature than the standard buffers such as Tris-HCl or HEPES[1]. DTT was added to prevent cysteine oxidation. The condensate sample contained ~0.0004% (w/v) microspheres. We used 0.2 µm microspheres for the temperature-dependent particle tracking and 1 µm microspheres for the passive microrheology with optical tweezers (pMOT) measurements.

**Passive microrheology with optical tweezers (pMOT).** pMOT experiments were performed following our previously published protocol[2]. Peptide-ssDNA condensate samples were placed on an 18 x 18 mm microscope coverslip and sandwiched with a 75 x 25 x 1 mm thick microscope glass slide using three layers of double-sided tape (Scotch 3M). Mineral oil was then injected into the chamber surrounding the sample from all directions to prevent sample evaporation with time. The sample was then loaded onto a correlative optical-tweezer and confocal microscopy setup (LUMICKS, Ctrap™). The sample was equilibrated for approximately 20 minutes until all droplets have settled on the coverslip surface and no fusion event was observed. By that time, several condensates were observed to contain 1-2 polystyrene beads. The optical trap was used to trap a bead within a condensate at minimal power (~10-50 µW). The trapped bead was then tracked using a brightfield camera at 500 Hz using a template-matching algorithm for 10-30 minutes. Approximately 20 condensates over three independent sample preparations were tested. Further details of pMOT experiments, including various control measurements, are reported in our earlier work[2].

**Temperature-controlled video particle tracking (VPT).** Peptide-ssDNA condensate samples were sandwiched between a coverslip and a glass slide similar to what is described in the pMOT section. The sandwich was then placed on a custom-built thermal stage (INSTEC) which is attached to a Zeiss primovert inverted microscope with a 100x oil-immersion objective lens. Teledyne FLIR blackfly S USB3 CMOS camera was used for imaging. Condensates were allowed to settle on the glass slide for approximately 20 minutes prior to the start of the measurement.

Each condensate contained about 20-50 fluorescent microspheres (200 nm). The temperature stage was first set to the lowest experimental temperature (5 °C). Due to the objective acting as a heat sink, the actual temperature within the sample was different from the preset temperature of the thermal stage. To obtain an accurate measurement of the sample temperature, we used a thermocouple with a heat insert that touches the glass directly inside the thermal stage chamber on the edge of the condensate sample. Within 10-20 minutes, the temperature reading of the thermocouple was observed to be equilibrating around 10 °C (preset temperature = 5 °C). To start imaging, the microscope objective was focused on a particular condensate. A time-lapse video was collected for 2000 frames at a rate of 10 frames per second (exposure time is set to 100 ms). For condensates that showed high viscosity and slow motion of the particles, a frame time of 200 ms was used. The temperature was increased in steps of 5 °C and the sample was left to equilibrate for 10 minutes at each temperature. Then a similar time-lapse video of the same condensate was collected to record the motion of the particles at the new temperature. This procedure was repeated to collect 5-7 points between 11 °C and 70 °C (preset temperature = 90 °C). Three trials were done for three independent sample preparations.

**Turbidity measurements.** Samples were prepared in a tube by mixing the peptide and the ssDNA at the desired mixing ratio and a fixed peptide concentration of 1 mg/ml. The buffer of these samples contains 25 mM MOPS (pH 7.5), 25 mM NaCl, and 20 mM DTT. The sample was then placed on a UV-Vis spectrophotometer (NanoDrop 1C) and the solution turbidity at 350 nm was measured for three independent samples. Before measuring the turbidity of peptide-ssDNA samples, the instrument was blanked using the experimental buffer. The values of the turbidity were averaged for each mixing ratio and the error was estimated as half the range of experimentally measured values.

**Fluorescence Recovery after Photobleaching (FRAP).** Samples were prepared at 5 mg/ml concentration of both peptide and ssDNA in a buffer containing 25 mM MOPS (pH 7.5), 25 mM NaCl, and 20 mM DTT. To measure peptide diffusion within $[RGRGG]_5$-ssDNA condensates, we added ~400 nM of the Alexa488-labeled $[RGRGG]_5$ peptides to the buffer before mixing the other components. Similarly, we used Alexa488-labeled $[RGYGG]_5$ peptides for $[RGYGG]_5$-ssDNA condensates. The final sample was then sandwiched between a glass slide and a coverslip using double-sided tape. Mineral oil was inserted into the chamber to surround the sample and prevent evaporation. The sample chamber was then placed on a confocal microscope (LUMICKS, CTrap™). For all FRAP experiments, we fixed the bleaching region to 0.6 x 0.6 μm. This was done to ensure that the changes in the FRAP recovery time are only due to the probe dynamics and not due to the larger bleaching area[3-5]. Further, we performed the FRAP experiments only on condensates with diameters that are significantly larger than 3 μm (at least 5x bleaching ROI radius) to rule out interfacial resistance effects[5]. For each sample, we measured the FRAP traces for 4-6 condensates. The recovery traces were corrected for photofading and normalized with respect to the bleaching depth following previously published procedures[6]. The recovery traces were then averaged, and the error was estimated as the standard deviation at each time point. To estimate the recovery half-time, we fitted individual recovery traces using the following equation [3, 6]

$$I_{recovery}(t) = \frac{I_0 + I_\infty \dfrac{t}{\tau_{1/2}}}{1 + \dfrac{t}{\tau_{\frac{1}{2}}}} \quad (1)$$

Here, $I_0$, $I_\infty$, and $\tau_{1/2}$ are fitting parameters, the latter represents the FRAP recovery half-time. The diffusion timescale was then calculated by dividing the recovery half-time by the square of the radius of the bleaching area ($\tau_D = \tau_{\frac{1}{2}}/R^2$). The reported FRAP diffusion timescale was taken as the average of the values extracted from the fits of individual FRAP measurements. The error was estimated as the standard deviation of the values.

**Data analysis: pMOT.** The output of the pMOT experiments is two-dimensional trajectories of trapped particles within condensates. These trajectories are analyzed to calculate the complex modulus of the condensate as a function of frequency $\omega$

$$G^*(\omega) = G'(\omega) + i\,G''(\omega) \tag{2}$$

Where $G'$ and $G''$ are the frequency-dependent elastic and viscous moduli, respectively. Briefly, the trajectories are first detrended using a spline-based detrending algorithm to remove the long-time drift of the trajectories. Next, each component (X and Y) of the trajectory is treated separately. For each one-dimensional trajectory (X or Y), we obtain the trap stiffness using the equipartition theorem[7-9]

$$\kappa_x = k_B T/\langle x^2\rangle \tag{3}$$

Where $\kappa_x$ is the optical trap stiffness in the $x$ direction, $T$ is the temperature in Kelvin, and $k_B$ is the Boltzmann constant. Next, a normalized position autocorrelation function $g(\tau)$ is calculated. The autocorrelation function value at $\tau = 0$ is extrapolated using a spline algorithm. The autocorrelation function is then transformed into the frequency domain $\hat{g}(\omega)$ using a discrete Fourier transform algorithm[7-9]

$$-\omega^2\hat{g}(\omega) = i\omega g(0) + \frac{\left(1 - e^{-i\omega t_1}\right)(g_1 - g(0))}{t_1} + \dot{g}(\infty)e^{-i\omega t_N} + \sum_{k=2}^{N}\left(\frac{g_k - g_{k-1}}{t_k - t_{k-1}}\right)\left(e^{-i\omega t_{k-1}} - e^{-i\omega t_k}\right) \tag{4}$$

Lastly, the complex modulus is calculated using[7-9]

$$G^*(\omega) = G'(\omega) + i\,G''(\omega) = \frac{\kappa}{6\pi a}\left(\frac{i\omega\hat{g}(\omega)}{1 - i\omega\hat{g}(\omega)}\right) \tag{5}$$

Where $a$ is the radius of the particle (0.5 μm). For each trial, we extract two sets of $G'$ and $G''$ values for each frequency. The total number of $G'$ and $G''$ sets is ~40 sets for ~20 condensates over three independent sample preparations. Finally, at each frequency, we average the values of $G'$ and $G''$ and report the final value. The error is calculated as the standard deviation. For all the systems tested, we exclude the top and bottom 5% of the values to eliminate any outliers. Further details of the experiment and data analysis, including various controls for the effects of condensate surface and glass surface on the measurement, are reported in our earlier work[2, 7].

**Data analysis: Temperature-controlled video particle tracking.** Movies of condensates containing diffusing polystyrene particles at different temperatures were processed first using Fiji-ImageJ software[10]. Each movie contained about 2000 frames. The particles were tracked using the TrackMate software plug-in in Fiji[11]. In particle tracking, intensity filters were used to exclude any particle aggregates from tracking. The extracted trajectories were corrected for drifting by subtracting the center of mass trajectory, which was calculated using the velocities of particles in the following way

$$\boldsymbol{X_{COM}}(t) = \boldsymbol{X_{COM}}(k\Delta t) = \boldsymbol{X_0} + \sum_{k=0}^{k\Delta t} \frac{1}{N} \sum_{i=1}^{N} \boldsymbol{v_i} \Delta t \tag{6}$$

Where $k$ is the frame number, $\Delta t$ is the frame time, and $N$ is the number of particles per frame. $\boldsymbol{X_0}$ is the center of the mass vector of the first frame. After correcting the trajectories for drifting, the mean squared displacement (MSD) was calculated as

$$MSD(m\Delta t) = \frac{1}{\Delta t} \frac{1}{N-m} \sum_{i=1}^{N-m} (x_{i+m} - x_i)^2 + (y_{i+m} - y_i)^2 \tag{7}$$

Where $m$ is the lag time in frames. The MSD was then fitted using[2, 12]

$$MSD(\tau) = 4D\tau^{\alpha} + N \tag{8}$$

To obtain the diffusion coefficient $D$ of the particles. For all the systems, we insured that the value of the diffusivity exponent $\alpha$ is equal to 1 by choosing an appropriately long frame time to ensure measuring the terminal viscous behavior. The diffusion coefficient is then converted to viscosity using the Stokes-Einstein equation[2, 12]

$$\eta = \frac{k_B T}{6\pi D R} \tag{9}$$

Where $R$ is the radius of the particles and $T$ is the temperature of the sample. This analysis was done for movies collected at different temperatures and the value of the viscosity was plotted against the temperature. To obtain the activation energy, we plotted $\ln \eta$ against $1/T$ and fitted the resulting data to Equation 2 in the main text. Each peptide-ssDNA condensate system was tested three times in three different sample preparation. The average value of the activation energy is reported, and the error is estimated as half the range of values. As a control, we measured the viscosity of the Water-Glycerol solution at 90% glycerol using our VPT-based approach and compared it with the published values in the literature[13, 14], which revealed good agreement (see Fig. S5).

### Supplementary Tables

| System | $E_A$ (RT) | $E_A$ (RT) | $E_A$ (RT) | $E_A \pm \Delta E_A$ (RT) |
|---|---|---|---|---|
| $[RPRPP]_5$-dT40 | 10.35 | 8.44 | 8.72 | 9 ± 1 |
| $[RGRGG]_5$-dT40 | 15.97 | 18.87 | 17.23 | 17 ± 1 |
| $[RGYGG]_5$-dT40 | 23.27 | 29.05 | 25.96 | 26 ± 3 |
| $[RGRGG]_5$-dT20 | 19.69 | 18.44 | 19.00 | 19.0 ± 0.6 |
| $[RGRGG]_5$-dT90 | 18.44 | 17.78 | 18.48 | 18.2 ± 0.4 |
| $[RGRGG]_5$-dT200 | 19.12 | 19.18 | 17.55 | 18.6 ± 0.8 |

**Table S1.** Activation energy values for all the peptide-ssDNA condensates tested in this study. Each sample was measured three times with independent sample preparations. The reported value is the average of the three measurements and the error is calculated as half the range of values. The units are $RT$ at T = 25 °C, where $R$ is the universal gas constant (1 $RT$ is 2.479 kJ/mol).

### Supplementary Figures

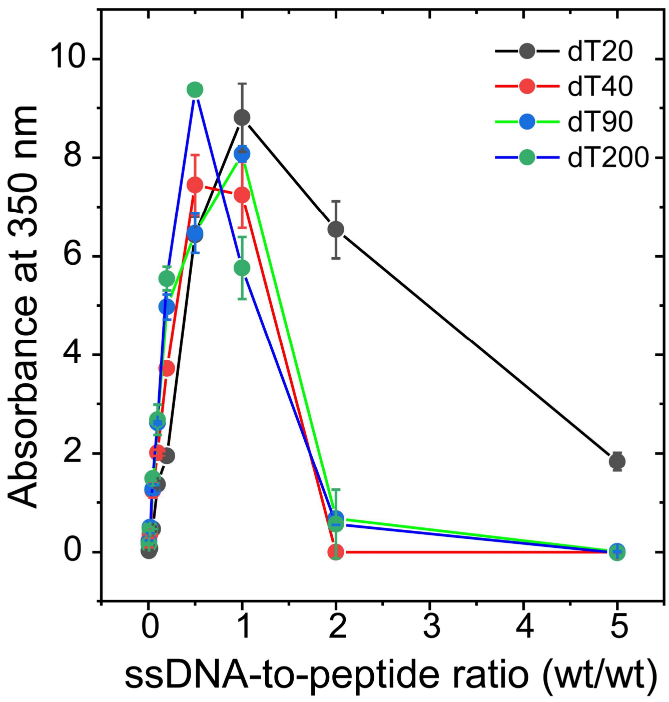


**Figure S1.** A plot showing absorbance at 350 nm (turbidity) of $[RGRGG]_5$-dT$n$ mixtures at different mixing stoichiometries. dT$n$ is a poly(dT) ssDNA with length $n$ equals 20, 40, 90, or 200.

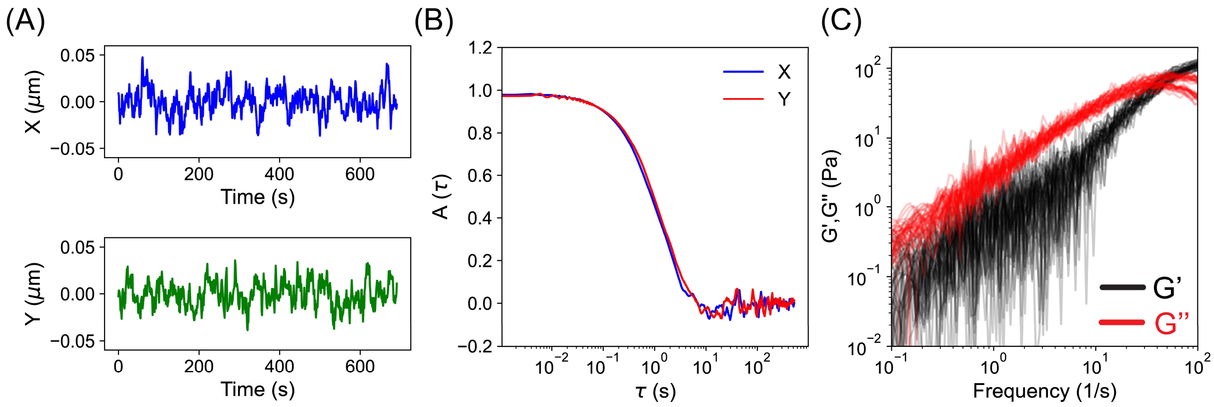


**Figure S2.** **(A)** Representative trajectories of a trapped bead within a $[RGRGG]_5$-dT40 condensate in the X and Y directions. **(B)** Position autocorrelation curves of the bead motion in the X and Y directions. **(C)** Viscoelastic moduli for twenty $[RGRGG]_5$-dT40 condensates. The reported data in Figure 1B in the main text is the average of these moduli.

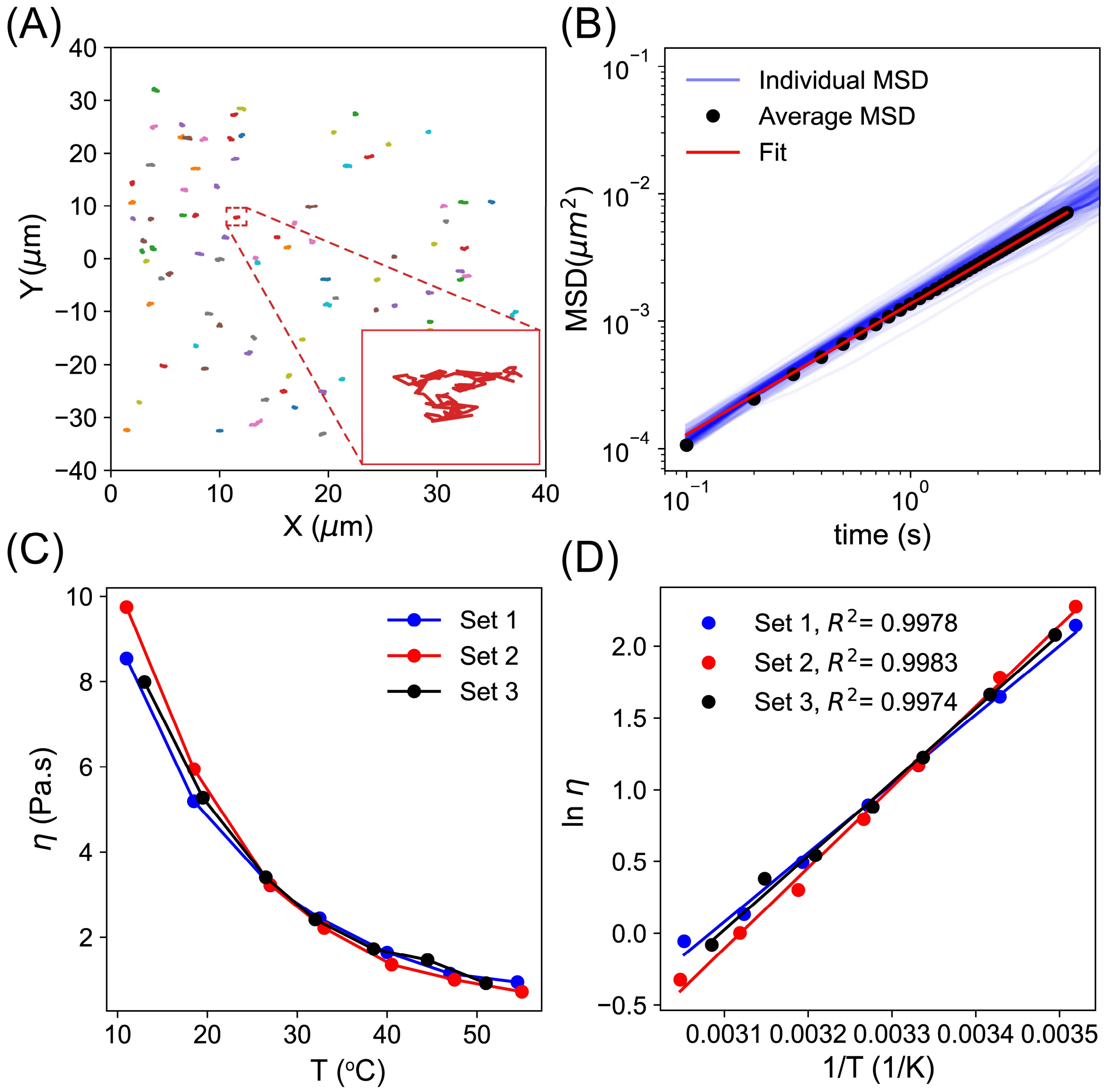


**Figure S3. (A)** Representative trajectories of 200 nm beads within a [RGRGG]$_5$-dT40 condensate as measured by video particle tracking at room temperature. **(B)** Individual MSD plots for all the beads within the condensate shown in (A). The black symbols are the ensemble-averaged MSD. The red line is a fit of the data according to Equation 8. **(C)** Viscosity variation with temperature for three independently prepared [RGRGG]$_5$-dT40 condensates. **(D)** Arrhenius plots for the data shown in (C) for the three trials. The lines are fits to the data using Equation 2 in the main text. The corresponding activation energy values are reported in Supplementary Table S1.

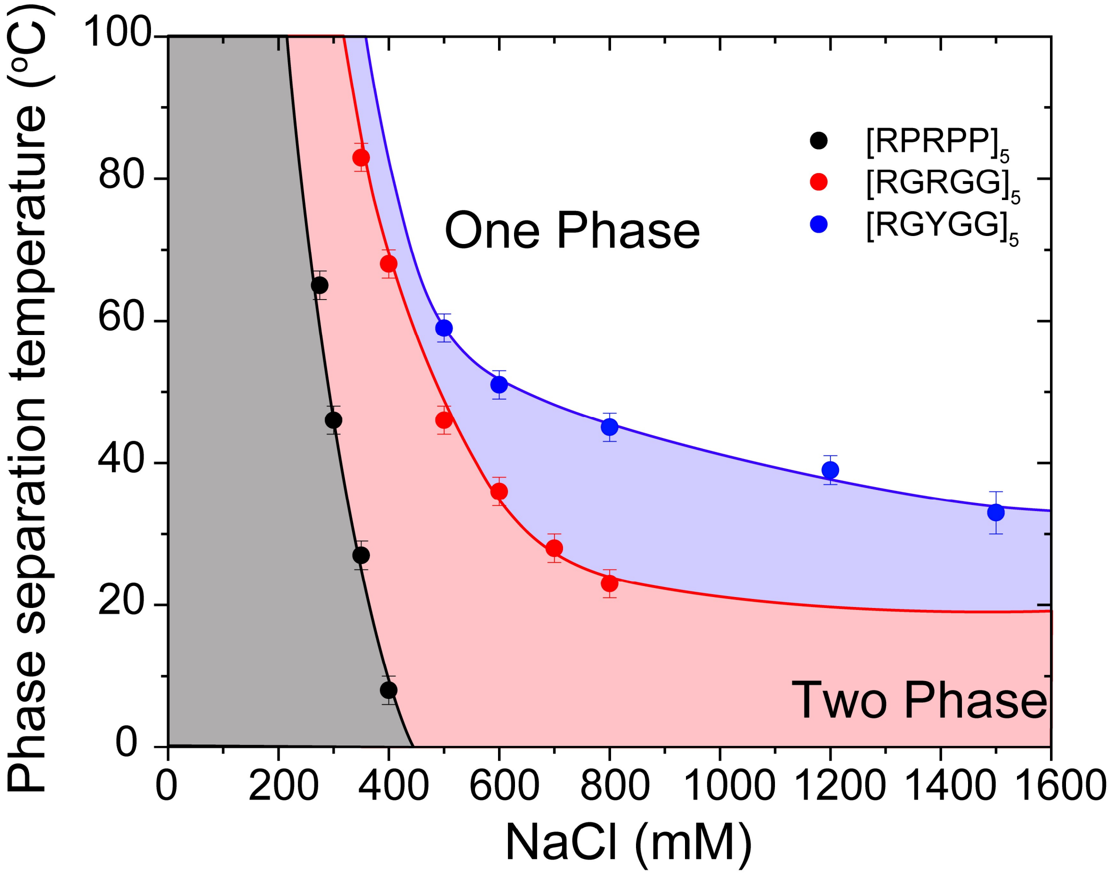


**Figure S4.** Temperature−salt phase diagrams for peptide-rU40 condensates. This data is repurposed from our previous work[2]. The shaded regions indicate temperature and salt conditions where phase separation occurs. All samples were prepared at 5 mg/ml polypeptide and 2.5 mg/ml rU40 RNA (analogous to dT40) in 25 mM Mops, 25 mM NaCl, and 20 mM DTT. Note that both $[RGRGG]_5$ and $[RPRPP]_5$ have similar stickers (Arg residues) even though they exhibit vastly different material properties as well as upper critical solution temperatures[2] (Fig. 2 in the main text). This can be attributed to the fact that spacers modulate peptide interactions with nucleic acids through the effective solvation volume and/or steric forces associated with the peptide-ssDNA complexes. Further, our observations that the flow activation energy decrease when mutating Gly spacers to Pro spacers further confirm that spacers affect the intermolecular interactions between the peptide and the ssDNA chain as well as the chain solvation properties. Accordingly, there are further nuances to the sticker-spacer classification of residues beyond what is considered in this work.

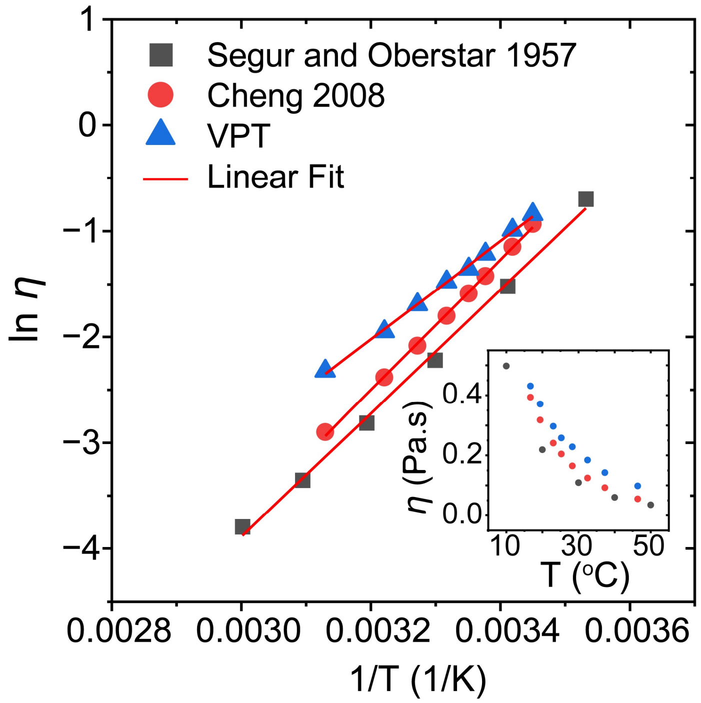


**Figure S5.** Arrhenius plots for the viscosity of Glycerol-Water mixtures as obtained from our VPT measurements (blue triangles; this study), a previous study by Segur and Oberstar[13] (black squares), and the viscosity formula from Cheng[14]. Red lines are linear fits according to Equation 2 in the main text. The obtained values of the activation energy are 15.6, 20.7, and 19.6 RT (38.7, 51.2, and 48.7 kJ/mol), respectively. The inset shows the viscosity value as a function of temperature for the same data sets.

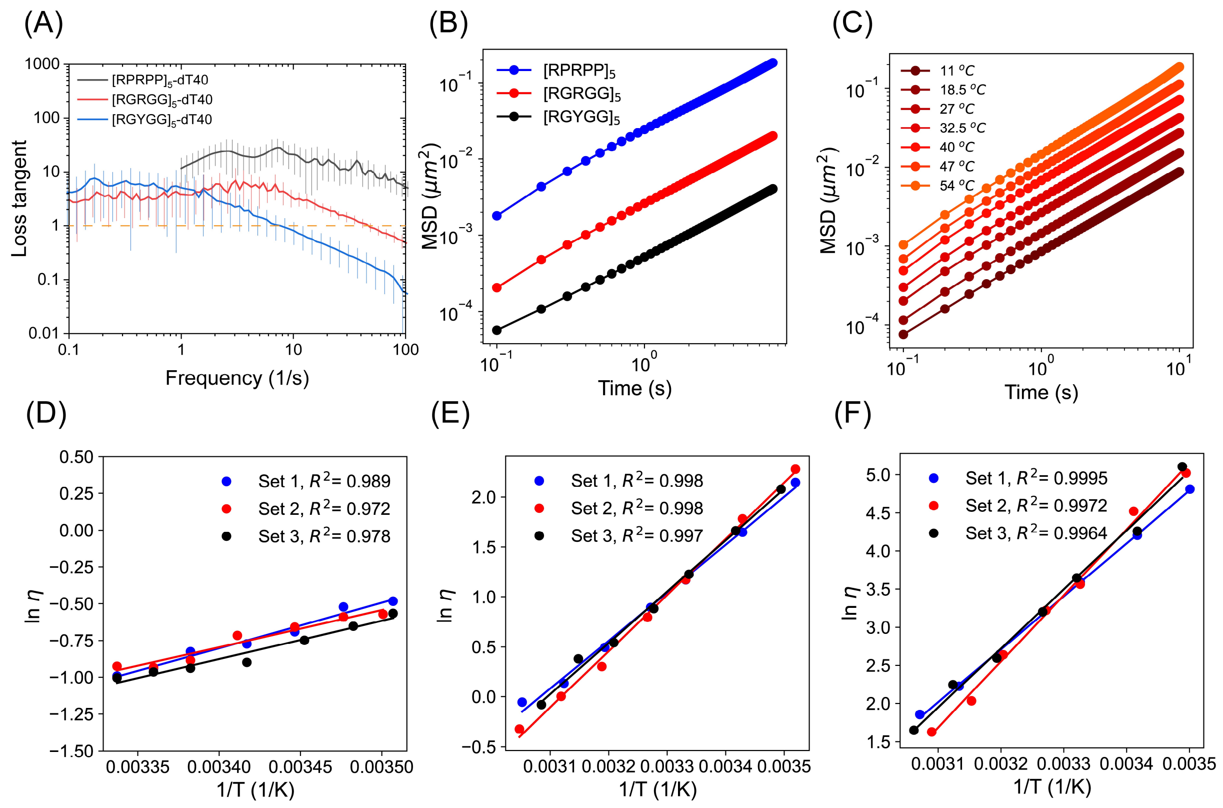


**Figure S6. (A)** Loss tangent ($G''/G'$) of peptide-ssDNA condensates showing distinct terminal relaxation times depending on the peptide sequence. **(B)** Ensemble-averaged MSD of 200 nm particles within peptide-dT40 condensates. The peptides shown here are [RGRGG]$_5$, [RPRPP]$_5$, and [RGYGG]$_5$. **(C)** Ensemble-averaged MSD of 200 nm particles within a [RGRGG]$_5$-dT40 condensate at different temperatures. **(D, E, F)** Arrhenius plots for three independently prepared peptide-dT40 condensates for the three peptides; [RPRPP]$_5$, [RGRGG]$_5$, and [RGYGG]$_5$, respectively. The lines are fits to the data using Equation 2 in the main text. The corresponding activation energy values are reported in Supplementary Table S1.

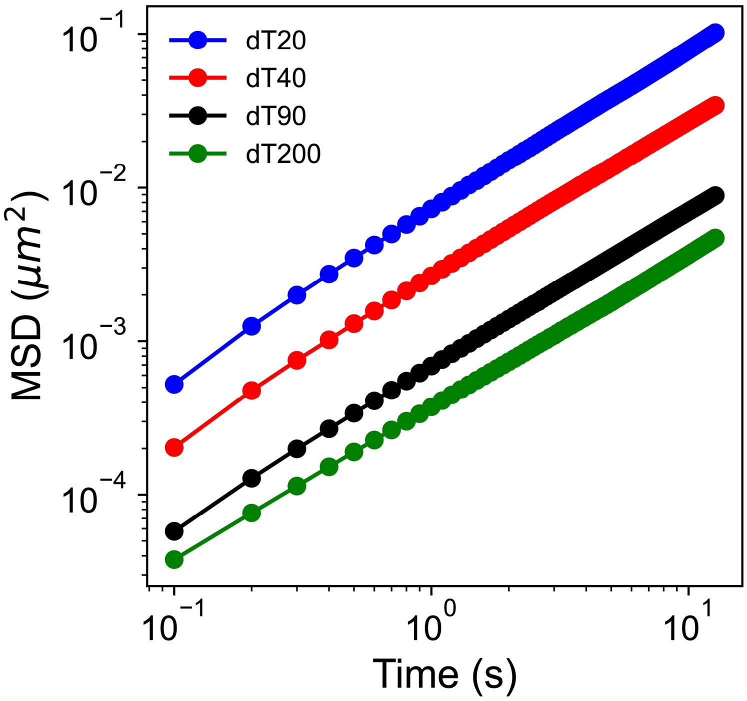


**Figure S7.** Ensemble averaged MSD of 200 nm particles within $[RGRGG]_5$-dT$n$ condensates where $n$ is the length of the ssDNA. Data are shown for $n$ = 20, 40, 90, and 200.

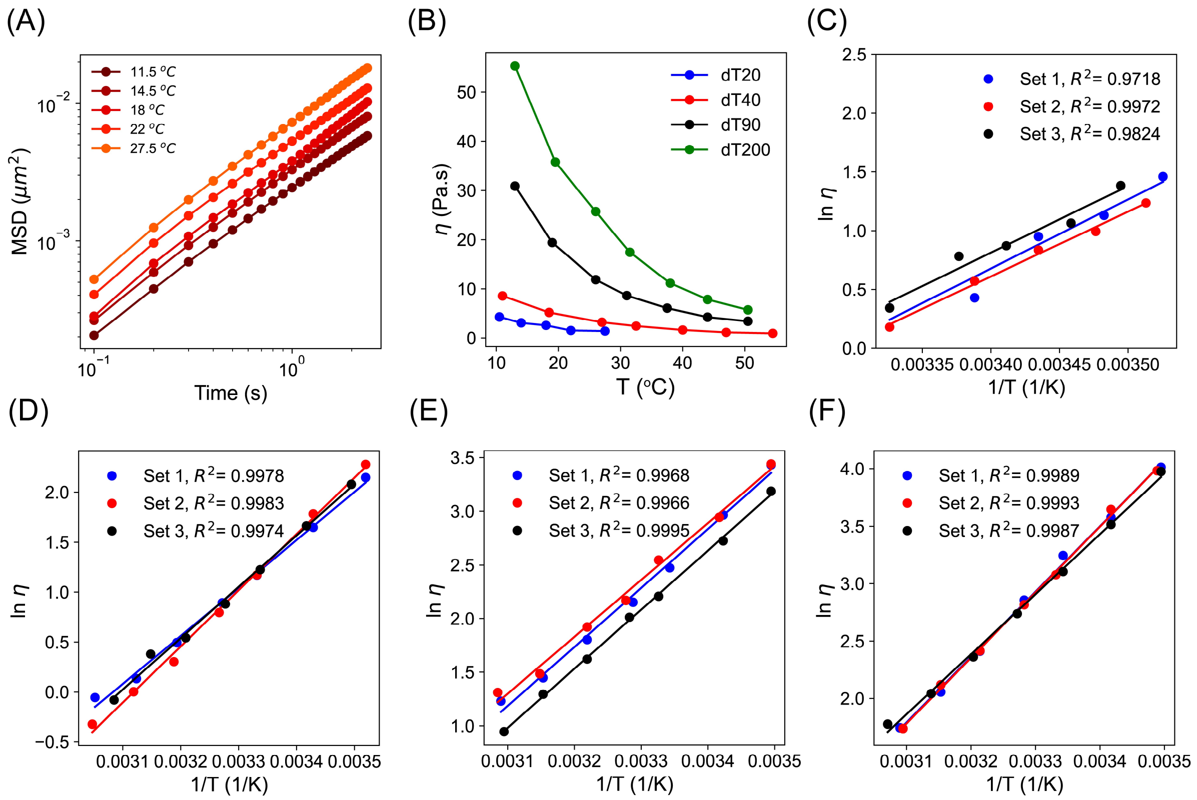


**Figure S8. (A)** Ensemble-averaged MSD of 200 nm particles within $[RGRGG]_5$-dT20 condensates at different temperatures. **(B)** Viscosity variation with temperature for $[RGRGG]_5$-dT$n$ condensates, where $n$ is the length of the ssDNA. The DNA lengths tested are 20, 40, 90, and 200. These are the same data as shown in Figure 4a in the main text. **(C-F)** Arrhenius plots for three independently prepared $[RGRGG]_5$-dT$n$ condensates where $n = 20$, 40, 90, and 200, respectively. The lines are fits to the data using Equation 2 in the main text. The corresponding activation energy values are reported in Supplementary Table S1.

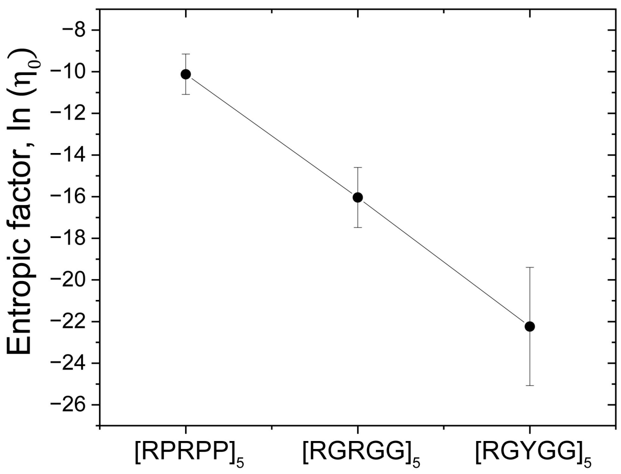


**Figure S9.** The extrapolated entropic factor of peptide-dT40 condensates from the data shown in Figure 2f in the main text. The entropic factors are calculated from the intercept of the linear fit according to Equation 2 in the main text.
